# Structural Characterization of Native RNA Polymerase II Transcription Complexes and Nucleosomes in *Drosophila melanogaster*

**DOI:** 10.1101/2025.02.03.636274

**Authors:** Natalie L. Venette-Smith, Rishi K. Vishwakarma, Varun Venkatakrishnan, Roberta Dollinger, Josie Schultz, Paul Babitzke, Ganesh Anand, David S. Gilmour, Jean-Paul Armache, Katsuhiko S. Murakami

## Abstract

Structural studies of eukaryotic RNA polymerase II (Pol II) transcription complexes often depend on *in vitro* assembly by mixing purified Pol II with synthetic DNA/RNA scaffolds, recombinant transcription factors, and/or histones, followed by stalling transcription at defined positions by adding selected nucleotide triphosphate substrates. These studies have yielded remarkable results for understanding nucleosome transcription by Pol II with elongation factors but may fail to represent transcription in native conditions. To investigate Pol II transcription within metazoan cells, we developed an approach to isolate the native transcription complexes from *Drosophila melanogaster* embryos. Utilizing one-step FLAG-tag affinity purification and mild chromatin treatment with Micrococcal Nuclease (MNase), we preserved the native transcription complex for cryo-EM and proteomics studies. *In silico* purification through the cryo-EM classifications determined structures of multiple forms of native transcription complex, nucleosome and other macromolecules. Remarkably, we determined the structures of metazoan Rpb4/Rpb7 stalk-less elongation complex as well as hexameric nucleosome lacking an H2A/H2B dimer, revealing that diverse elongation complexes and nucleosomes are involved in active transcription *in vivo*. Nucleosome is positioned only downstream of Pol II in the nucleosome elongation complex, underscoring it as a major energy barrier and a time-consuming step during Pol II progression through nucleosomal DNA. Proteomics identified co-purified factors responsible for initiation and elongation stages of transcription, as well as RNA modification factors. This study lays the groundwork for structural study of native transcription in eukaryotes, with future work focused on studies of transient and minor populations of transcription complexes.

## Introduction

In all eukaryotes except for plants, three DNA-dependent RNA polymerases can be found in the nucleus: RNA polymerase I, II and III, abbreviated as Pol I, II and III, respectively. Pol II is responsible for transcribing messenger RNA (mRNA), long noncoding RNA (lncRNA) and a number of small RNAs (1). While Pol II alone has the capacity to synthesize RNA from a DNA template, binding of various factors allows it to proceed through different stages of transcription (initiation, elongation and termination), acquire diverse functions and provide opportunities for regulation.

Transcription initiation involves the assembly of Pol II with general transcription factors (GTFs) and the Mediator complex, followed by CDK7-mediated phosphorylation of the Serine-5 residues in the C-terminal domain (CTD) heptapeptide repeats of Pol II (2). Binding the Pol II with negative elongation factor (NELF) and Spt4/Spt5 (as known as DSIF) establishes promoter proximal pausing, a post initiation regulatory sub-step. Active transcription begins after the removal of NELF from Pol II, and bindings of elongation factors such as TFIIS, elongation factor 1 (ELOF1; aka Elf1), Spt6 and the PAF complex (3). Furthermore, histone chaperones such as facilitates chromatin transcription (FACT) to enhance Pol II transcription through the nucleosome DNA. When Pol II reaches the poly-A termination sequence, a series of factors terminate transcription by releasing RNA and Pol II from DNA.

Structural biology has been one of the major ways of probing the transcriptional machinery. It reveals how different factors interact with Pol II to elucidate their functions. The first high-resolution structure of Pol II was obtained by X-ray crystallography in 2000 (4). Since then, over 200 structures of Pol II complexes have been determined by X-ray crystallography or cryo-electron microscopy (cryo-EM) (5). Currently, cryo-EM is the primary method for structural biology, as it does not require crystallization, does not require a large quantity of sample, and allows visualization of heterogeneity and conformational dynamics of targeted macromolecule. This makes it well-suited for determining the structures of transcriptional complexes containing multiple factors as well as elusive intermediates during nucleosomal DNA transcription (6, 7).

In many of these studies, the transcription complexes are assembled *in vitro* with purified Pol II, synthetic DNA/RNA as well as recombinant transcription factors and/or histones, and some cases, adding limited nucleotide triphosphate (NTP) substrate to move Pol II on DNA template at defined locations. Recent studies of the nucleosome elongation complex (EC) have provided snapshots of Pol II moving through nucleosomal DNA, showing disassembly of a nucleosome at downstream DNA followed by reassembly at upstream DNA, either in the absence (8, 9) or in the presence of the histone chaperone (10). However, the insights from these studies are derived from experimental conditions specifically designed to capture transient intermediates, leaving it uncertain whether they faithfully represent native transcription processes within cells. Recently, cryo-electron tomography (cryo-ET) in combination with cryo-Focused Ion-beam Milling (cryo-FIB) is employed to visualize thinned flash-frozen biological samples in their near-native environment (11). This *in situ* approach allows for atomic-resolution structural characterization of gigantic macromolecules such as ribosomes (−4.5 MDa molecular weight) as they can be easily located in crowded cellular tomograms followed by averaging in 3D to improve resolution (12). The technique is in continuous development, however, and since Pol II (−500 kDa molecular weight) is −9 times smaller than ribosome, identifying their positions and orientations within tomograms remains a technical issue. This leads to the question: How does one characterize native Pol II transcription complexes?

To address this gap, we establish a method for visualizing endogenous transcription complexes directly isolated from *Drosophila melanogaster* (Drosophila) embryos. We used large population of CRISPR/Cas9-modified Drosophila containing a double FLAG-tag on Rbp1-CTD, to obtain sufficient Pol II complexes for cryo-EM and proteomics studies. Although the affinity purified Pol II complexes remain heterogenous and contain impurities, *in silico* purification of targeted macromolecules through cryo-EM data processing allowed us to determine multiple forms of transcriptional complexes, nucleosomes and other macromolecules co-purified with Pol II.

## RESULTS

### Isolation of native Pol II transcription complexes from FLAG-tagged Rpb1 Drosophila embryos

In this study, we used a double FLAG-tagged Rpb1 Drosophila strain for isolating native Pol II transcription complex (13). Using populations of −10,000 adults, we collected −75 g of 0-12 hr embryos for nuclei preparation (**Fig. 1A**). Next, we used MNase, an endo- and exo-nuclease that preferentially targets the linker DNA between nucleosomes (14), to digest chromatin while preserving nucleosome bound with Pol II and other factors. MNase digestion at 0.5 U/µL for 10 mins at room temperature produced DNA fragments of −150 bp and −250 bp, corresponding to mono- and di-nucleosomes, respectively (**Fig. 1B**). Nuclei lysis followed by removing insoluble materials, Pol II transcription complex was isolated by using one-step FLAG-affinity purification. The presence of Pol II in elution fractions was confirmed by colloidal Coomassie-stained SDS-PAGE analysis, which showed well-defined bands of Rpb1, Rbp2 and Rpb3 subunits, with impurities (**Fig. 1C**). Western blot analysis confirmed presence of Rpb3 and histone H3 in the elution fractions (**Fig. 1D**). Quality and quantity of the Pol II transcription complex were estimated by negative stain EM.

**Figure 1.**
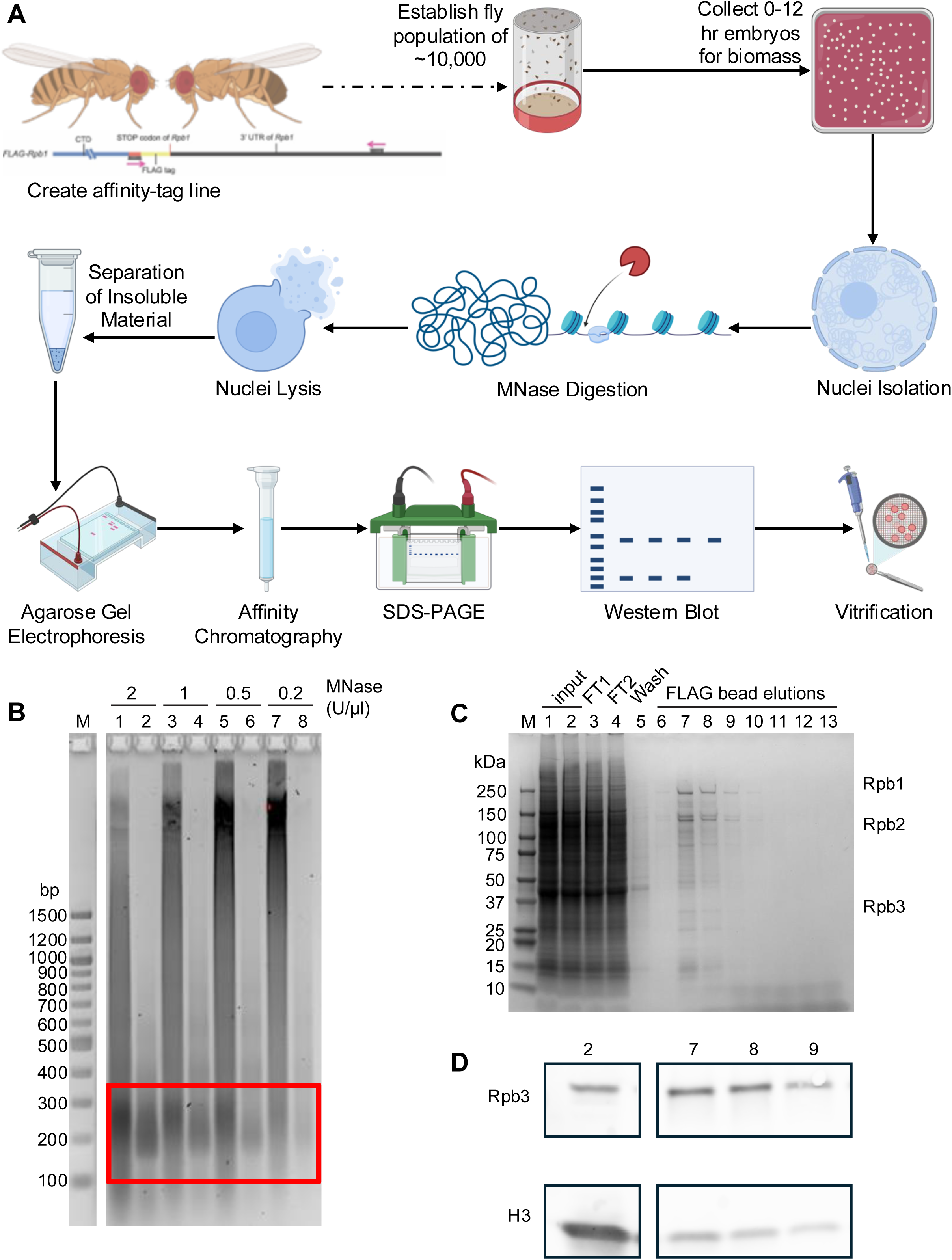
Methodology and purification of native Pol II transcription complex from Drosophila embryo. **(A)** Diagram showing the processes of Drosophila embryo collection, MNase treatment and nuclear extract preparation, purification of Pol II transcription complexes and characterization, and vitrification for cryo-EM data collection. **(B)** Chromatin DNA digested by MNase. DNA fragments from mono- and di-nucleosomes are indicated in red box. Lanes: (1,2) 2 U/µL MNase digestion, (3,4) 1 U/µL MNase digestion; (5,6) 0.5 U/µL MNase digestion, (7,8) 0.2 U/µL MNase digestion, (1,3,5,7) insoluble fractions after MNase digestion, (2,4,6,8) soluble fractions after MNase digestion. **(C)** Coommassie stained SDS-PAGE protein gel of affinity-purified samples. Lanes: (1,2) input, (3) Sepharose bead flow-through, (4) FLAG bead flow-through, (5) FLAG bead wash, (6–13) FLAG bead elution. Subunits of Pol II are indicted. **(D)** Western blot detection of Pol II by anti-Rpb3 antibody (top) and nucleosomes by anti-histone H3 antibody (bottom). Lanes are labeled as in panel **(C)**.

### In silico purification of native Pol II and other macromolecules isolated from Drosophila embryos

Cryo-EM data were collected using a Talos Arctica 200 kV microscope equipped with a Falcon 4 detector (**SI Appendix Table 1**). Despite the heterogeneity of the affinity-purified Pol II complex containing several impurities (**Fig. 1C, SI Appendix Figs 1A**), computational particle picking and 2D/3D classifications during cryo-EM data processing enabled us to enrich particles belonging to each targeted macromolecule, a process we term “in silico purification.” From 30,103 micrographs, we extracted −15 million particles, which underwent multiple rounds of 2D and 3D classifications, yielding subsets of macromolecules (**Fig. 2; SI Appendix Figs. 1-2**). These subsets include Pol II, nucleosome, a putative U1 spliceosome, and an octahedral density attributed to the dihydrolipoyllysine-residue succinyltransferase (DLST) component of the 2-oxoglutarate dehydrogenase complex. Further 3D classification separated 876,764 Pol II particles into four major subunits including: 1) EC with stalk; 2) EC without stalk; 3) nucleic acid-free apo-form and 4) nucleosome EC.

**Figure 2.**
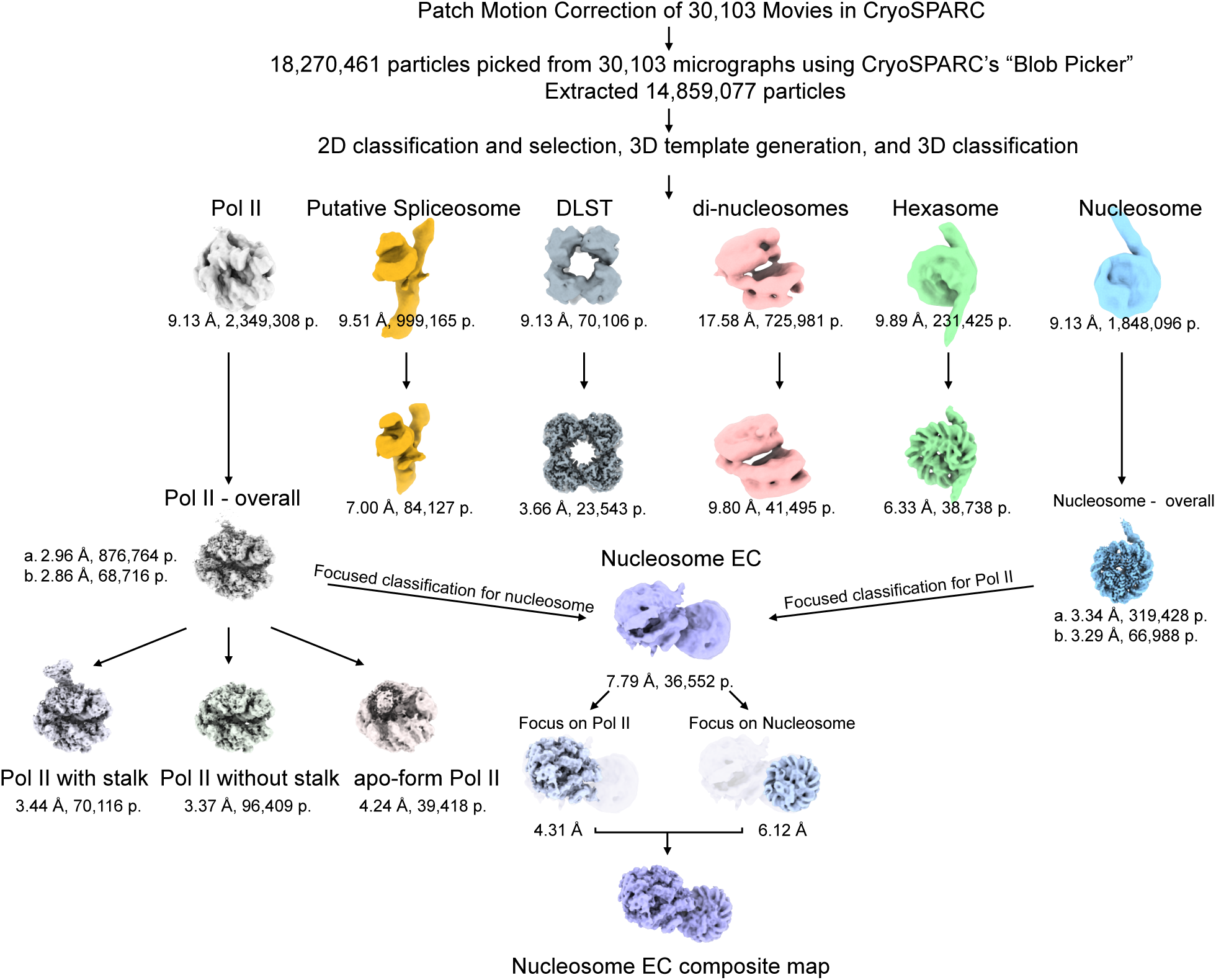
In silico purification of the Pol II transcription complexes and nucleosomes. A simplified scheme of *in silico* purification of the macromolecules by cryo-EM data processing (for details, see ***Materials and Methods***). The preprocessed macromolecule particles were subjected to 3D classification, yielding six unique subsets, representing Pol II (gray), putative U1 spliceosome (orange), DLST (dark gray), stacked nucleosomes (pink), hexasome (green) and nucleosome (blue). The first row represents the initial subsets that were further processed to obtain the final reconstructions (the second row). Pol II particles were classified further into three subsets, containing: EC with Rpb4/7 stalk, EC without Rpb4/7 stalk, and apo-form Pol II. Further individual analysis of Pol II and nucleosome subsets yielded enriched particles of nucleosome EC (middle, violet). Pol II and nucleosome within the nucleosome EC were refined by focused/local refinement, and combine these maps to generate a composite map of nucleosome EC.

**Table 1.**
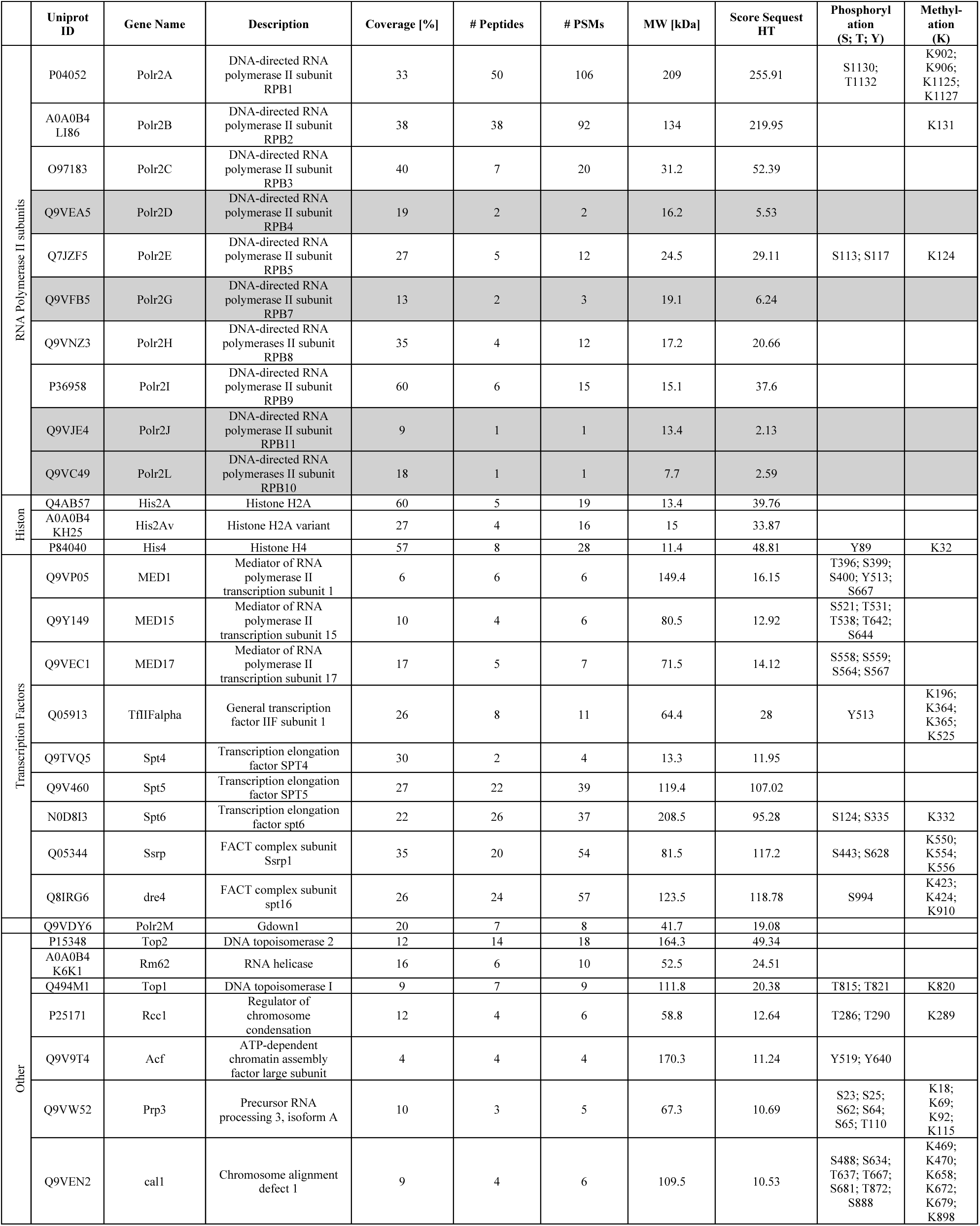
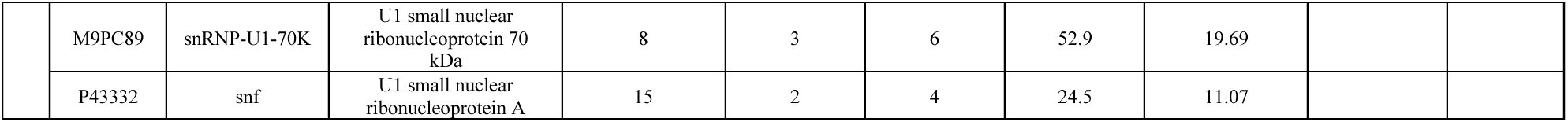
Proteins identified in purified endogenous Drosophila Pol II through liquid chromatography tandem mass spectrometry. In gray, Pol II subunits below the selected Score Sequest HT threshold of 10.

### Cryo-EM structural analysis of the native Drosophila Pol II elongation complex

The cryo-EM map of the apo-form Pol II was solved at a lower resolution (4.24 A) and was visibly anisotropic (**SI Appendix Figs. 1C and 3A**) due to preferred particle orientation. Thus, we could not reliably construct its model. In contrast, the EC with and without the Rpb4/7 stalk show well-defined isotropic cryo-EM densities (**Fig. 3A, SI Appendix Figs. 1D and E, SI Appendix Table 2, SI Movie 1**). The cryo-EM map shows Pol II, 10 bases of DNA/RNA hybrid and 25 bases of double-stranded downstream DNA. The 3’ end of the RNA is in the post-translocated site (i site) and nucleotide binding site (i+1 site) is vacant (**Fig. 3B**). The EC shows no sign of RNA backtracking (15), nor a tilted state of the DNA/RNA hybrid (16). The cryo-EM density for DNA/RNA hybrid is strong and traceable until 10 bases from the 3’ end of RNA, which provides the first direct evidence showing that the native Pol II EC accommodates 10 bases of DNA/RNA hybrid. The lid, rudder (Rpb1) and flap (Rpb2) form the RNA exit channel to guide nascent RNA toward outside of Pol II but cryo-EM density of single-strand RNA beyond DNA/RNA hybrid is not resolved (**Fig. 3B**). The density corresponding to upstream DNA is weak but traceable, however, single-stranded non-template DNA within a transcription bubble is not visible (**Fig. 3A**). Since DNA and RNA sequences within the EC are mixed due to *in situ* Pol II transcription at different locations in genome, we modeled DNA and RNA with the following sequences: template DNA as poly(dT); non-template DNA as poly(dA), and the nascent RNA as poly(A). The overall structure of Pol II is nearly identical to the structures of other eukaryotic Pol II, including those from yeast (17, 18) and mammal (19, 20) (**Fig. 3C**).

**Figure 3.**
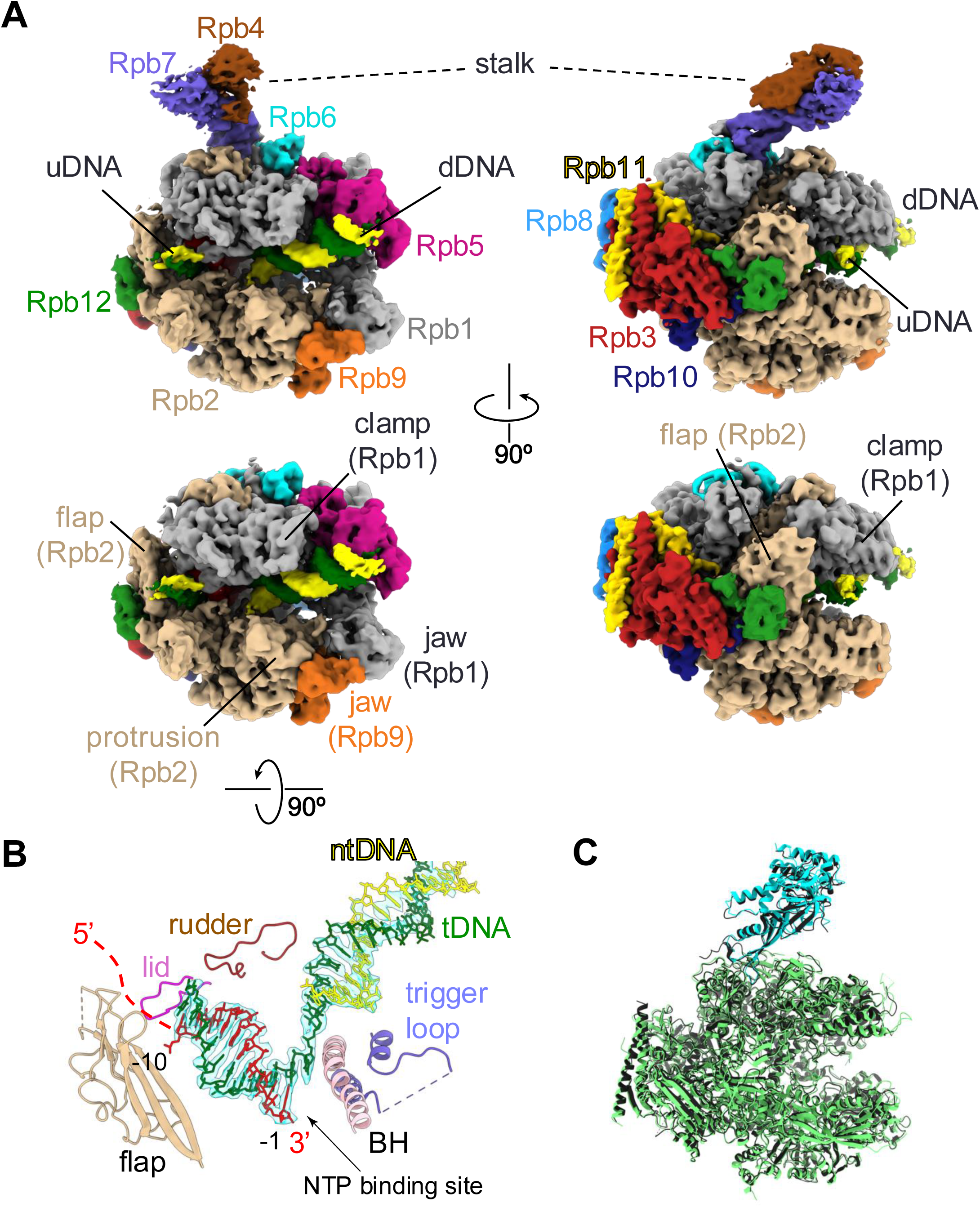
Cryo-EM structure of the Pol II EC. **(A)** Orthogonal views of the cryo-EM density map of the Drosophila EC with stalk (top) and without stalk (bottom). Pol II subunits (Rpb1-12), DNA and RNA are colored and labeled (dDNA, downstream DNA; uDNA, upstream DNA). Several Pol II domains are also indicated. **(B)** Cryo-EM density map (transparent cyan) of DNA and RNA in the EC is shown with DNA/RNA (stick model, tDNA, template DNA; ntDNA, non-template DNA) and motifs of Pol II (cartoon model, BH, bridge helix) in the refined EC. NTP binding site of the active site is indicated. **(C)** Comparison of the structures of the Pol II within Drosophila EC (green, main body; cyan, stalk) and mammalian EC (black) (PDB: 6GML, (16)). Structures were superimposed via main body of Pol II with an RMSD of 0.795 A.

The cryo-EM density of the Rpb4/Rpb7 stalk of Pol II was less well-defined compare with the main body of Pol II (see Pol II-overall in **Fig. 2, SI Appendix Fig. 1B**), suggesting its mobility and/or sub-stoichiometric occupancy. Focused 3D variability analysis (3DVA) in cryoSPARC revealed two subsets in which stalk was either fully present (EC with stalk, **Fig. 3A** top; **SI Appendix Fig. 1D**) and completely absent (EC without stalk, **Fig. 3A** bottom; **SI Appendix Fig. 1E**), and these cryo-EM maps are determined as overall resolutions of 3.44 and 3.37 A, respectively. Not only the EC with stalk, but also EC without stalk show strong densities of DNA and RNA, indicating that the stalk-less Pol II is actively participated in transcription within cells. The numbers of particles corresponding to the EC with and without the stalk are about the same (**Fig. 2; SI Appendix Table 2**) indicating that a significant proportion of the stalk-less Pol II participates in gene expression in metazoan cells.

### Cryo-EM analysis of the native *Drosophila* nucleosome

The in silico purification also enriched a large pool of nucleosome particles and yielded subsets including: 1) octameric nucleosome (called “nucleosome” throughout this manuscript); 2) hexameric nucleosome (hexasome) and 3) di-nucleosome (**Fig. 2**). We obtained a 3.29 A reconstruction of a nucleosome (**SI Appendix Fig. 2A; SI Appendix Table 3**), which shows an asymmetric arrangement of entry and exit sites DNA, where one site is −11 bases shorter compared to the other (**Figs. 4 A,B**). Throughout the manuscript, we refer to the shorter and longer DNA strands outside from nucleosome core as the entry and exit sites, respectively. This high-resolution reconstruction of nucleosome visualizes features such as side-chain densities and the DNA phosphate backbone (**SI Appendix Fig. 4A**). As these nucleosomes contain a mix of native DNA sequences, the base assignment was not possible, thus we modeled them using a 601 Widom sequence (21). These nucleosomes were very similar in structure to recombinant octameric nucleosomes (22, 23).

**Figure 4.**
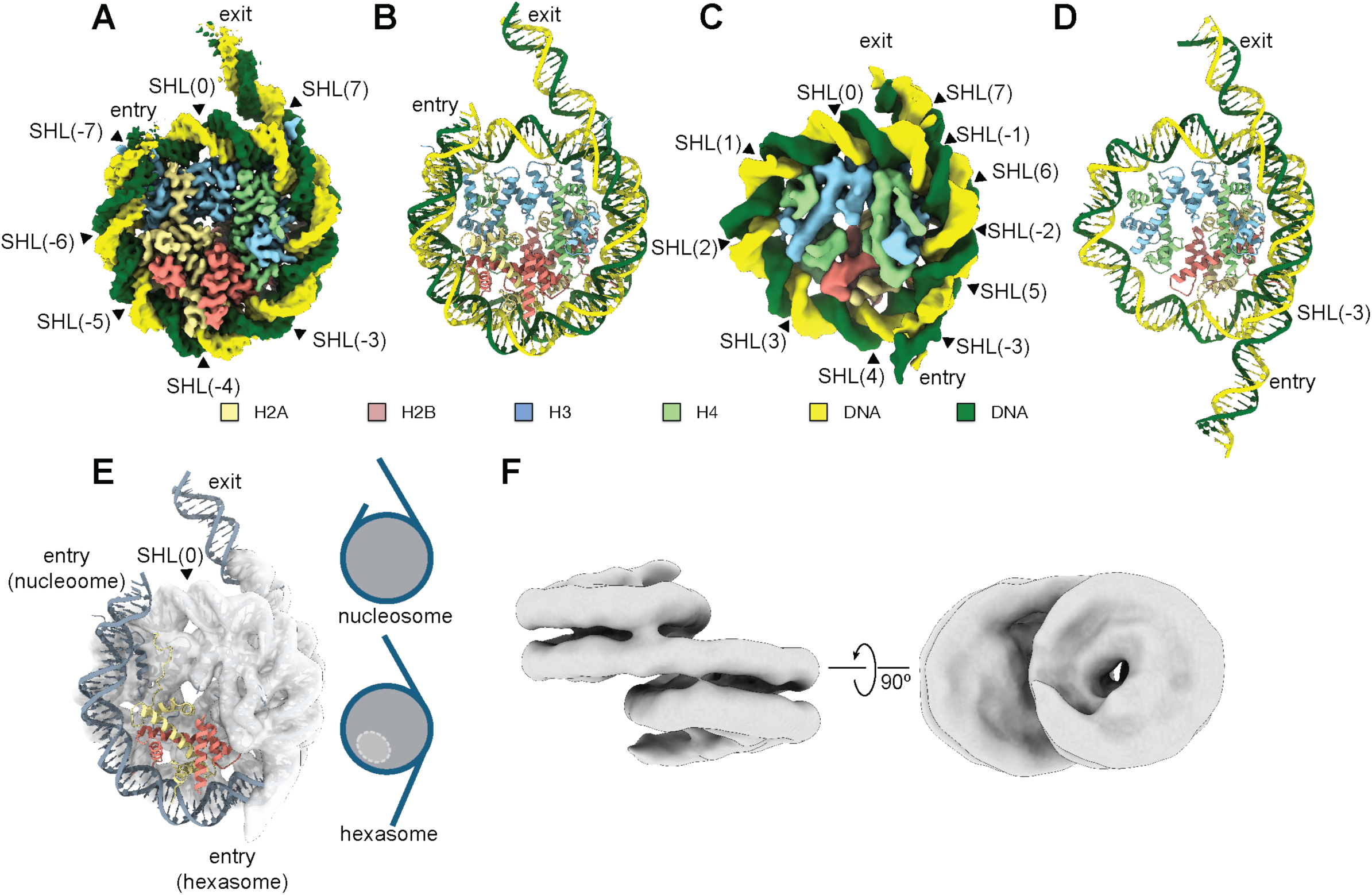
Cryo-EM structures of nucleosome, hexasome and di-nucleosome. Cryo-EM density maps of Drosophila nucleosome **(A)** and hexasome **(C)**. Super Helical Location (SHL) are marked starting from SHL(0) at the dyad axis of nucleosome. Refined models of the nucleosome **(B)** and hexasome **(D)** are shown using the color scheme from (A) and (C). **(E) (left)** A nucleosome model (dark blue) is superimposed in the cryo-EM density map of the hexasome. A H2A/H2B dimer of nucleosome, which is missing in hexasome, is indicated and colored (light yellow and light red); (**right**) Comparative schematic representation of the nucleosome and hexasome. Positions of the entry and exit sites DNA and SHL are labeled. **(F)** Cryo-EM density map of an overall reconstruction of the di-nucleosomes.

We also obtained a 6.33 A reconstruction of a hexasome (**Figs. 4C,D; SI Appendix Fig. 2B; SI Appendix Table 3**), which lacks a single copy of the H2A/H2B histone dimer as compared with nucleosome (**Fig. 4E**). The quality of the hexasome reconstruction allowed for unambiguous positioning of histone subunits and the DNA path around the hexasome core. Approximately 100 base pairs of DNA wrapping around the hexasome (**Fig. 4D**), which is −50 base pairs shorter than the −150 base pairs of DNA wrapping around the nucleosome (**Fig. 4B**). In the hexasome, the DNA contacting with histone core starts at the superhelical location SHL(−3), continues through SHL(0), and ends at SHL(7) at the exit site. Due to the absence of one H2A/H2B dimer in the hexasome, the DNA prior to the SHL(−3) site unwraps from histone core, thus the DNA at the entry and exit sites point in opposite directions (**Fig. 4E**). These observations are consistent with previous characterizations of hexasome by structural and genome wide nucleosome profiling (24–27).

Intriguingly, we also obtained a subset representing di-nucleosome, in which two nucleosomes are partially stacked in a stair-wise manner, while also being connected by the linker DNA (**Fig. 4F; SI Appendix Fig. 2C**). A similar arrangement of di-nucleosome in recombinant experiments has been previously observed through X-ray crystallography (28) and cryo-EM (29) and was also reported in complex with a chromatin remodeling enzyme, CHD1 (26), wherein the two nucleosomes were bridged by the remodeler.

### Cryo-EM structure of Drosophila Nucleosome Elongation Complex

After confirming the presence of both Pol II and nucleosome in the elution fractions of FLAG-affinity purification (**Fig. 1D**) and through the cryo-EM data processing to obtain reconstructions of the Pol II and nucleosome, we employed further *in silico* purification from these subsets to enrich the nucleosome EC particles (**Fig. 2**). In particular, careful analyses of the Pol II and nucleosome 2D classes revealed some showing extra density beyond their well-defined densities. Using particles belonging to these 2D classes, we created an ab initio cryo-EM map as a reference of the nucleosome EC followed by carrying out the 3D classification of particles belonging to Pol II and nucleosome to enrich particles belonging to nucleosome EC. The isolated nucleosome EC was well-defined and could be refined to an overall resolution of 7.79 A, limited by the Pol II and nucleosomes moving each other. To address this, we refined them independently (focused/local refinement), resulting in a stalk-less Pol II and nucleosome at resolutions of 4.31 A and 6.12 A, respectively **(SI Appendix Fig. 3F).** We combined these maps to create a composite map of nucleosome EC for a model building and refinement (**Fig. 5; SI Appendix Table 2**). The cryo-EM map clearly shows template DNA and endogenous RNA in the Pol II and downstream DNA extending from the DNA binding channel of Pol II wraps around histones in nucleosome. The downstream nucleosome has clear DNA features such as sugar-phosphate backbones, and major and minor grooves, as well as an identifiable the histone core (**SI Movie 2**). Notably, the nucleosome EC is observed in one state only, with Pol II located at the nucleosome position SHL(− 5). About 20 bases DNA unwrapped from the nucleosome for Pol II moving toward it, but maintain the DNA interacting with H2A/H2B. The jaw (Rpb1), protrusion (Rpb2) and jaw (Rpb9) domains of Pol II face to nucleosome but there is no physical contact between Pol II and nucleosome (**Fig. 5C**). The cryo-EM structure of native nucleosome EC determined in this study, which is derived from Pol II transcriptions starting from promoters, is similar to one of previously reported nucleosome EC structures that were assembled and prepared in vitro (8) (PDB: 6A5P, **SI Appendix Figs. 3 D, E)**. It is likely that other previously reported states of the *in vitro* reconstituted nucleosome ECs are transient and represent minor states during the Pol II transcription through nucleosome DNA *in vivo*.

**Figure 5.**
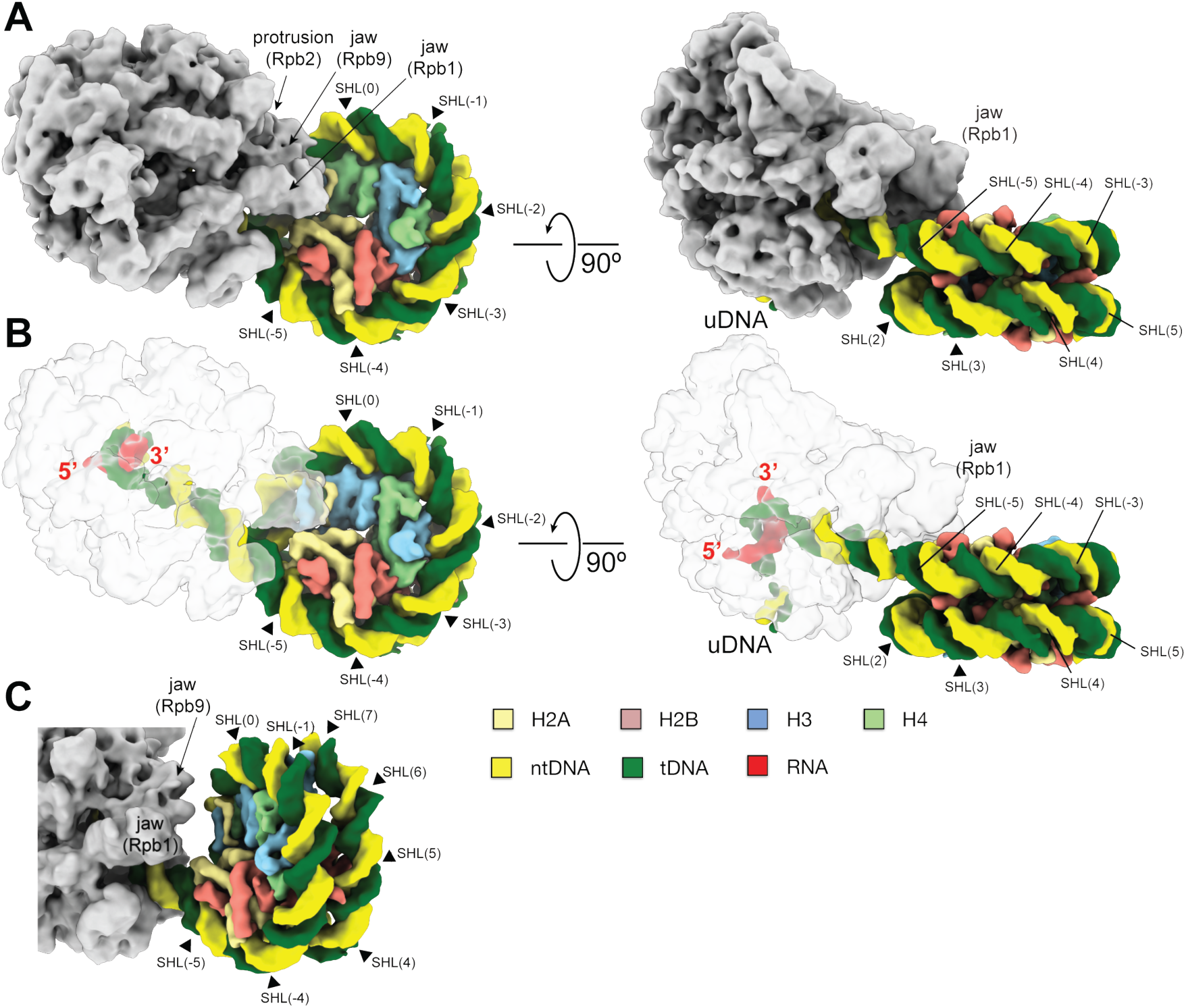
Cryo-EM structure of the native nucleosome EC. **(A,B)** Orthogonal views of the cryo-EM densities of the Drosophila nucleosome EC paused at SHL(−5) position of nuclesomal DNA. Pol II (gray), DNA, RNA and histones are colored. **(B)** Orthogonal views of the cryo-EM densities of nucleosome EC highlighting the nucleosome core, DNA and RNA with transparent Pol II density (white). (**C**) A magnified view of the interface between Pol II and nucleosome. Domains of Poll II and SHL positions are labeled.

### Mass spectrometry identification of proteins co-purified with Pol II

To identify factors co-purified with Pol II, we analyzed them by mass spectroscopy. Trypsin digested samples were analyzed by bottom-up liquid chromatograph-tandem mass spectrometry (LC-MS/MS), a technique commonly applied to identify proteins from macromolecular assemblies (30). Peptide search against the entire *D. melanogaster* genome was used for protein identification with searches for post-translational modifications (PTM) such as phosphorylation on Serine, Threonine and Tyrosine, and lysine methylation. We identified a total of 28 proteins participating to the Pol II transcription, nucleosome formation, chromatin remodeling and RNA processing, and other proteins of interest with a Sequest HT Score greater than 10 (**Table 1**).

While 10 of the 12 subunits of Pol II could be identified, only Rpb 1, 2, 3, 5, 8, 9 subunits met the cut-off criteria of Score Sequest HT greater than 10 (**Table 1**). Nucleosomes were confirmed due to the presence of histones H2A, H2A variant, and H4. Three subunits of mediator (MED 1, 15, 17) and subunit 1 of TFIIF were identified, as well as DSIF (including Spt4 and Spt5 subunits) and metazoan-specific Pol II binding factor Gdown1 were identified (31). Although its score was just below the arbitrary cut-off, the presence of NELF was also confirmed (**SI Dataset 1**). The proteomics study suggests that an endogenous promoter-proximal pausing complex including EC, DSIF and NELF can possibly be isolated and enriched by the affinity and *in silico* purifications for future cryo-EM characterization. Most factors identified by mass spectrometry are involved in transcriptional elongation underscoring the most Pol II participate in the active RNA synthesis. The histone chaperone FACT (Ssrp and Spt16) (10, 32) was also identified, although it was not found in the cryo-EM sturdy of nucleosome EC (**Fig. 2**). Other proteins of interest are DNA topoisomerase 1 and 2, RNA processing factors, chromatin/chromosome factors and the U1 snRNP spliceosome (U1 snRNP and snRNP A subunits) (**Fig. 2, SI Appendix Fig. 2D**; **Table 1**).

## DISCUSSION

In this study, we used a FLAG-tagged Rpb1-CTD of Pol II to isolate native transcriptional complex from Drosophila embryos. The high-yield Drosophila population system allowed us to harvest sufficient embryos for isolating endogenous Pol II complex for cryo-EM and proteomics. In silico purification facilitated the isolation, enrichment, and construction of 3D cryo-EM maps of individual macromolecules, achieving separations not possible with other methods. For examples, distinguishing Pol II with or without the stalk, and nucleosome or hexasome, are challenging using chromatography purification due to their chemical and molecular weight similarities. While affinity tagging at specific site can achieve some separations, in silico purification offers a more straightforward approach without requiring genome editing. This method is powerful and convenient for studying heterogeneous and dynamic transcription processes in a cellular context. In addition, all transcription complexes characterized here originated from native promoters, enabling the visualization of endogenous, rather than artificially assembled, complexes.

The Rpb4-Rpb7 heterodimeric stalk binds around the RNA exit channel of Pol II. Rpb7 is essential in budding yeast *S. cerevisiae*, but Rpb4 is only essential under stress conditions and during starvation for retaining viability (33, 34). Rpb4 is essential in fission yeast *S. pombe* (35), *Drosophila* (36) and for zebrafish development (37). The stalk functions in closing the DNA binding clamp (17, 18), providing binding platform of numerous transcription factors (38–41) and for forming Pol II dimer (42, 43). The stalk also functions in RNA transport and translation in the cytoplasm (44). In *S. cerevisiae*, several studies suggested that the stalk can dissociate from the EC (45), but the stalk separation from the transcribing Pol II remains controversial in other studies (46). One challenge was to observe the stalk-less Pol II actively participating transcription *in vivo*. In this study, *in silico* purification allowed us to isolate it (**Fig. 2**), giving us a direct evidence of the stalk-less Pol II involved in the active transcription. This finding aligns with chromosomal immunostaining of *Drosophila* third instar larvae, showing that not all transcriptionally-active Pol II co-localize with Rpb4, as seen on polytene chromosome (36).

Numerous transcription factors associate with Pol II in each stage of transcription (38, 39). The cryo-EM analysis of *in vitro* reconstituted transcription complexes revealed how these factors influence the Pol II activity and providing functions to Pol II for carrying out stage specific activities. However, we did not observe Pol II particles unambiguously associated with any factors, despite their identification by proteomics study. Through further data processing, we identified small subsets of EC showing extra densities around the Spt5 and Spt6 binding sites (**SI Appendix Figs. 3 F,G**). However, they resulted in low resolution reconstructions and thus could not be unequivocally identified. Perhaps, most Pol II engaging in expressing transcription during rapid cell cycle embryogenesis in Drosophila do not require elongation factors (Spt4/5, Spt6) and/or histone chaperones (e.g., FACT) due to lower densities of nucleosomes at highly expressing genes (47) and coupling between DNA replication and transcription (48).

We also isolated native metazoan nucleosomes including nucleosome, hexasome, and di-nucleosome (**Fig. 4**). Nucleosomes co-eluted with Pol II from FLAG-affinity column may result from their abundance in the nuclear extract or association with apo-form Pol II, as shown previously (49). Additionally, we co-purified spliceosomes and DLST, a mitochondrial enzyme. Spliceosomes may associate with Pol II to couple transcription and splicing (50). DLST, an enzyme localized in mitochondria, may be isolated because of contamination. However, it was reported that a small percentage of DLST remains in the nucleus to provide succinyl-CoA to histone acetyl/succinyltransferase, KAT2A, a component of the eukaryotic transcription coactivator Spt-Ada-Gcn5-Acetyltransferase (SAGA) (51).

Affinity-purified Pol II from Drosophila embryos enabled cryo-EM and proteomic studies, revealing diverse Pol II and nucleosome complexes actively engaged in transcription. Our findings likely represent the most stable and abundant transcription complexes due to the lack of cross-linking prior cryo-EM specimen preparation. Crossing FLAG-tagged Rpb1 lines with those encoding tags for GTFs, elongation factors, or histones could enrich minor Pol II complexes for future studies. CRISPR/Cas9 could further expand this system for isolating diverse complexes.

The Drosophila transcription system has uncovered key mechanisms in eukaryotic gene regulation, such as promoter-proximal pausing (52). While differences in transcriptional activity between Drosophila and humans exist, the similarities in transcription factors and the advantages of the Drosophila genetic system make it a powerful model for studying transcription. This approach can also be extended to other developmental stages, offering a versatile model for structural and proteomic analyses.

### Experimental Procedures

#### Growth and collection of *Drosophila* embryos

A *D. melanogaster* strain (Bloomington Drosophila Stock Center number 99015) expressing Rpb1 with a double FLAG-tag at the C-terminus was used in this study (53). The sequences expressing the double FLAG-tagged Rpb1 were introduced into the endogenous Rpb1 gene in Lu and Gilmour (13). Drosophila populations of −10,000 adults were housed in a 30 cm x 60 cm plastic cylinder pipe for 10 days. Starting on day 3, 0-12 hr embryos were collected on grape juice/agar plates and stored for a maximum 3 days at 4 °C. Embryos were strained through 2 layers of mesh to remove any adult flies, washed with water, dechorionated by mixing with 50% bleach for 30 sec and washed with deionized water before being stored at −80 °C. These steps were repeated until enough embryos were collected for the subsequent experiments.

#### Generation of nuclear extract

We prepared nuclear extract using a published procedure with modifications (54). 75 g of partially thawed, dechorionated embryos were resuspended in 120 mL Buffer A (1 M sucrose, 10 mM HEPES-HCl (pH = 7.5), 4 mM MgCl_2_, 0.1 mM EGTA, 1 mM B-mecaptoethanol (BME), 1 mM sodium bisulfite, 0.2 mM PMSF, 1 mM benzamidine HCl, and protease inhibitors (1.6 µg/mL benzamidine-HCl, 1 µg/mL aprotinin, 1 µg/mL pepstatin A, and 2 µg/mL leupeptin), homogenized with Potter Elvehjem tissue homogenizer, strained through a layer of MiraCloth, divided evenly into 6 Sorvall tubes, and centrifuged at 1,200 rpm at 4 °C for 10 mins to pellet debris. Supernatant was transferred to fresh Sorvall tubes and centrifuged at 6,000 rpm at 4 °C for 20 mins. The supernatant was aspirated off the nuclei pellets, taking care to remove all the fatty layer. Nuclei were resuspended in 5 mL buffer A, and dounced in a 7 mL homogenizer with first a loose pestle then a tighter pestle. Homogenized nuclei were poured over 10 mL of high sucrose buffer (1.75 M sucrose, 10 mM HEPES-HCl (pH = 7.5), 2 mM MgCl_2_, 0.1 mM EGTA, 1 mM BME, and protease inhibitors), mixed gently to generate a rough sucrose gradient and prevent nuclei aggregation at the interphase, and centrifuged at 12,500 rpm at 4 °C for 30 mins. The supernatant was aspirated off the nuclei pellets, taking care to remove all the fatty layer. Nuclei were resuspended in 1.5 mL of nuclear resuspension buffer (0.3 M sucrose, 10 mM HEPES-HCl (pH = 7.5), 2 mM MgCl_2_, 0.1 mM EGTA, 0.1 M NaCl, 1 mM DTT, 2 mM CaCl_2_, and protease inhibitors) and combined in a 15 mL Falcon tube before undergoing MNase Digestion.

Chromatin digestion was performed with 0.5 U/µL concentration MNase (#LS004797, Worthington Biochemical Corporation) at room temperature for 10 mins. As per the manufacturer’s definition, a single unit of MNase is the amount of enzyme necessary to change the optical density at 260 nm by 1.0 when incubated at 37 °C, pH 8.0 and DNA used as substrate. The reaction was quenched by adding EGTA to a final concentration of 15 mM and placed on ice. Triton X-100 was added for a final concentration of 0.5 % and incubated on ice for 10 mins before nuclei were mechanically lysed by passing sample through a 21-gauge needle 4 times. Lysed nuclei were gently mixed at 4 °C for 20 mins and then centrifuged at 16,000x g at 4 °C for 10 mins to separate insoluble and soluble material. The supernatant was transferred into clean 1.5 mL microfuge tubes, flash frozen in liquid nitrogen and then stored at −80 °C.

#### Purification of RNA Polymerase II Complex

Nuclear lysate was thawed and centrifuged at 16,000x g at 4 °C for 10 mins to remove any additional insoluble material before overnight dialysis at 4 °C into FLAG purification buffer (10 mM HEPES-HCl (pH = 7.5), 150 mM NaCl, 5% glycerol, 1 mM EDTA). Post-dialysis nuclear lysate was centrifuged at 16,000x g at 4 °C for 10 mins again to remove any remaining insoluble material. Protease inhibitors and 1/1000^th^ Proteoloc Protease Inhibitor (100X Expedeon Proteoloc Protease Inhibitor Cocktail - Expedeon 42516) were added directly to sample before affinity purification. All steps were performed at 4 °C, unless stated otherwise.

Sepharose 4B resin (Sigma-Aldrich 4B1200-100ML) at a ratio of 100 µL slurry per 1 mL nuclear lysate was prepared with two 5 min washes with FLAG purification buffer (supplemented with 1/1000x concentration protease inhibitors) before batch binding with sample for 10 mins. Supernatant was separated by centrifugation at 2,500 x g at 4 °C for 2.5 mins. Supernatant was transferred to anti-FLAG resin (Sigma-Aldrich ANTI-FLAG M2 Affinity Gel #A2220) at a ratio of 60 µL slurry per 1 mL nuclear lysate, which was prepared as described above, and gently mixed for batch binding for 3 hours. Resin slurry was transferred to a column and allowed to settle before the flow-through passed. The column was washed with FLAG purification buffer (supplemented with 1/1000^th^ protease inhibitors) with a 5 min incubation period. Pol II complexes were eluted from the column using elution buffer (10 mM HEPES-HCl (pH = 7.5), 150 mM NaCl, 5% glycerol, 1 mM EDTA, 350 µg/mL 3x FLAG peptide, 1/1000^th^ protease inhibitor) in 1 column volume (CV) fractions. Elution buffer was incubated on column before collection: fractions 1-2 at 30 mins, fractions 3-4 at 20 mins and fractions 5-8 at 10 mins.

SDS-PAGE followed by Western blot analysis using rabbit anti-Rbp3 (serum made in-house as described in Lu and Gilmour (53)) and rabbit anti-histone H3 (abcam #AB1791) antibodies confirmed presence of both Pol II and nucleosomes in the elution fractions. Alexa-488 fluorescent secondary antibodies (Invitrogen #A11008) and Amersham Typhoon Fluorescent Image Analyzer (GE Healthcare Bio-sciences) were used for Western blot analysis. Elution fractions containing both Pol II and nucleosomes were combined, dialyzed overnight in Flag Purification Buffer to remove any remaining 3x Flag Peptide, and concentrated by filtration (Amicon Ultra 0.5 mL 10000 NMWL UF501096). Qualitative assessment of the Pol II transcription complex was determined by negative stain (0.75% Uranyl Formate) before proceeding with vitrification.

#### Cryo-electron microscopy data collection and in silico purification of targeted macromolecules

Cryo-EM grids were prepared as follows. Quantifoil Au 2/2 grids (200 mesh, 2-µm hole size) were glow-discharged for 10 s at 15 mA in a PelCo easiGlow system (PelCo). 3.5 µl of the sample (1.8 mg/mL) was applied on the grid surface, and the grids were blotted and plunge-frozen in liquid ethane maintained at liquid-nitrogen temperatures, using a Vitrobot IV (FEI) maintained at 4 °C and 100% humidity. The data was recorded on a Talos Arctica G2 system at Penn State Huck Cryo-EM Core Facility, operated at 200 kV in nanoprobe mode, using a Falcon 4 camera at a nominal magnification of 150,000x corresponding to a calibrated physical pixel size of 0.944 A. The collection resulted in 30,103 images, each fractionated into 42 frames, accumulating 50 e^-^ per A^2^ total dose (0.0825 s per frame, 1.25 e^-^/A^2^/frame).

Raw movies were processed by cryoSPARC version 4.4.1 (55). We used *“Patch Motion Correction”*, followed by *“Patch CTF Estimation (Multi)”* to preprocess the images. 18,270,461 particles were picked by *“Blob picker”* and extracted 14,859,077 particles in a 440×440 box Fourier-cropped to 96×96. *“2D classification”* into 250 classes revealed multiple forms of macromolecules as seen on the raw images (**SI Appendix Fig. 1A**). The 2D classes representing Pol II, nucleosomes, spliceosome and DLST were selected independently and used to generate initial 3D templates using *“Ab initio”* reconstruction. We also selected 2D classes representing ice and other contaminants and used them to generate *ab initio* 3D templates attracting particles of impurities. These initial 3D templates were used to enrich particles into independent subsets (*in silico* purification) by using *“Heterogeneous Refinement”.* We then optimized the quality and resolution of the individual subsets and quantified their heterogeneity using a combination of *“Non-Uniform Refinement”* (56), *“2D classification”*, *“Heterogeneous Refinement”* and *“3D Variability”*. All final reconstructions were calculated using particles in box 440×440 (0.944 A/pixel) Fourier cropped to 360×360 (corresponding to 1.1538 A/pixel), following a gold-standard refinement by *“Non-Uniform Refinement”* and are reported at Fourier Shell Correlation = 0.143 (57) (**SI Appendix Table 2,3**).

##### Refinement and classification of Pol II particle subset

After refining the initial Pol II particles, we obtained an anisotropic reconstruction due to particle orientation bias. Using “*Heterogeneous Refinement”* in combination with *“2D classification”* and selection, we reduced bias and generated an isotropic reconstruction. Further optimization through signal-based selection using CryoSieve (58) yielded the final resolution of 2.86 A. Upon careful examination of the data, we found that the Rpb4/7 stalk was not well-resolved compare with Pol II containing 10 subunits. We performed a series of global and local classifications using “*Heterogeneous Refinement”* and *“3D Variance”*, resulting in three distinct Pol II reconstructions including: 1) apo-form, 2) EC with the stalk and 3) EC without the stalk. The individual subsets of EC with and without stalk were further refined, yielding 3.44 A and 3.37 A final resolution reconstructions, respectively (**Fig 2; Appendix Figs. 1B,D,E; Appendix Table 2**).

##### Refinement and classification of octameric nucleosome particles

We refined the octameric nucleosome subset using *“Non-Uniform Refinement”*. The reconstruction indicated that one side of the nucleosomal DNA, when displayed at different thresholds, disappears earlier than the other side. To investigate this further, we used *“Heterogeneous Refinement”* to classify the nucleosome particles into two subsets, varying in DNA length on one side. The highest quality nucleosome reconstruction was obtained in C1 symmetry at 3.34 A, which was then subjected to signal-based selection using CryoSieve, yielding a reconstruction at 3.29 A (**Fig. 2; SI Appendix Fig. 2A; SI Appendix Table 3**).

##### Refinement and classification of hexameric nucleosome particles

Hexameric nucleosomes were refined using *“Homogeneous Refinement”*, and the corresponding particles were reextracted at 1.1538 A/pixel and re-refined using *“Non-Uniform Refinement”*. The data was further processed and homogenized using *“Heterogeneous Refinement”*. The final reconstruction was refined to 6.33 A using *“Non-Uniform Refinement”* (**Fig 2; SI Appendix Fig. 2B; SI Appendix Table 3**).

##### Refinement and classification of stacked nucleosome particles

Stacked nucleosomes were refined using *“Homogeneous Refinement”*, and the corresponding particles were reextracted at 1.1538 A/pixel and re-refined using *“Non-Uniform Refinement”*. The data was further processed and homogenized using selection through 2D classification and *“Heterogeneous Refinement”*. The final reconstruction was obtained at 9.80 A using *“Non-Uniform Refinement”* (**Fig 2; SI Appendix Fig. 2C**).

##### Refinement and classification of nucleosome EC particles

A subset representing nucleosome EC was generated through iterative 2D classification and selection for enriching nucleosome EC particles from the Pol II and nucleosome subsets. We then used cryoSPARC’s *“Ab initio”* reconstruction to generate a 3D template, which was used *“Heterogeneous Refinement”* of particles from the Pol II and nucleosome subsets. Resulting particles were further analyzed using 2D classification to eliminate impurities, such as ice, and refined using *“Homogeneous Refinement”*. We then re-extracted the particles at 1.1538 A/pixel and refined the data using *“Non-Uniform Refinement”* to obtain the final overall reconstruction of 7.79 A. In this map, Pol II and nucleosome were not defined to high quality; therefore, we refined them individually in focused masks using *“Local Refine”*, resulting in the final reconstructions at 4.31 and 6.12 A, respectively. For modeling and refinement of the nucleosome EC structure, we combined these maps to generate a composite map using Phenix (**Fig 2; SI Appendix Fig. 1F; SI Appendix Table 2**).

##### Refinement and classification of putative spliceosome particles

Particles corresponding to the putative spliceosome in our data were refined using *“Homogeneous Refinement”*, reextracted at 1.1538 A/pixel, and re-refined. We performed extensive 2D classification analysis, which further confirmed that this subset suffered from a severe orientation bias, which drastically limited the quality of the final reconstruction (**Fig. 2; SI Appendix Fig. 2D**). CryoSPARC reported the final resolution using particles at 1.1538 A/pixel at 7.00 A, however the features of this reconstruction do not reflect this quality.

##### Refinement and classification of DLST particles

Throughout our processing, particles exhibiting a potential in-plane 4-fold -symmetry were found using 2D classification in every reported subset. We selected and pooled all the particles together. We then ran 2D classification on all these particles to further eliminate any contamination. Selected particles were used to generate ab initio 3D templates. The refinement against a selected 3D Coulomb potential density was first run in C1-symmetry. We then reextracted the particles at 1.1538 A/pixel and re-refined the data using *“Non-Uniform Refinement”* in C1, C2, C4, D4 and O symmetries. The octahedral symmetry yielded a reconstruction at 3.66 A, with clear secondary structure features (**Fig. 2; SI Appendix Figs. 2E, 5; SI Appendix Table 3**). To identify the macromolecule, we used ModelAngelo (59) using the *Drosophila* proteome database (Uniprot: UP000000803), which suggested that our Coulomb potential density map represented dihydrolipoyllysine-residue succinyltransferase component of mitochondrial 2-oxoglutarate dehydrogenase complex. To further confirm the assignment, we used AlphaFold, as well as an existing structure from *H. sapiens* (PDB: 6H05, (60)).

#### Model building

##### Pol II EC

A model of the Pol II EC was constructed by assembling AlphaFold 2-modeled (61) Pol II subunits (from Rpb1 to 12) using *Sus scrofa* Pol II (PDB: 6GML, (16)) as a guide. EC subunits were manually fitted into the cryo-EM density map using UCSF ChimeraX v1.8 (62), followed by real-space refinements using Coot (63) and Phenix (*phenix.real_space_refine*)(64). Modeling errors were corrected by DAQ refine (65) (**SI Appendix Tables 2, 4**).

##### Nucleosomes and hexasomes

The model of the *Drosophila* octameric nucleosomes were constructed using histone protein models from AlphaFold 2 with the 601 Widom DNA sequence used as a reference (PDB: 3LZ0, (23)) which was extended and adjusted to the Coulomb potential density map. This model was then used to construct the hexameric nucleosome. Both models were then refined using *phenix.real_space_refine* **(SI Appendix Tables 3,4).**

##### Nucleosome EC

To model the nucleosome EC, we manually placed the *Drosophila* EC and octameric nucleosome models into the composite map of nucleosome EC describe above. Using Coot, we further optimized the local fit and connected the DNA. We then fixed the model using *phenix.real_space_refine* **(SI Appendix Tables 2, 4**).

##### DLST

The model for DLST was constructed into the 3.66 A Coulomb potential density map. First, we used an AlphaFold 2 model and rigid-body fit it into one of the subunits. This monomeric model was then adjusted to the density and used to create a 24-subunit model. This model was refined through *phenix.real_space_refine* using “*Non-crystallographic symmetry*” restraints (**SI Appendix Tables 3, 4; SI Appendix Fig. 4B,C**).

#### Protein Identification through Mass Spectrometry

The protein identity was confirmed by the Huck Proteomics and Mass Spectrometry Core Facility (RRID: SCR_024462). The purified endogenous Pol II complex sample was prepared by reduction-alkylation of cysteines followed by a standard trypsin digestion (MS-grade trypsin, Pierce, Thermo Fisher Scientific, USA, MAN0011821). The sample was then subjected to liquid chromatography tandem mass spectrometry (LC-MS/MS) in an Orbitrap Eclipse mass spectrometer (Thermo Fisher Scientific). Data analysis was performed using Proteome Discoverer v2.5 against the *Drosophila* proteome (Uniprot: UP000000803) with default settings using Sequest HT search with fixed modifications set to carbamidomethylation of Cysteines and oxidation of Methionine and posttranslational modifications set to Serine, Threonine and Tyrosine phosphorylation, and Lysine methylation.

## Author Contributions

Fly embryo preparation (NLVS, RD, JS, DSG), Pol II preparation (NLVS, RD, JS, DSG), proteomics experiment (VV, GA), cryo-EM data collection and processing (NLVS, RKV, JPA and KSM), model building and refinement (JPA and KSM), manuscript writing (NLVS, RKV, VV, RD, PB, GA, DSG, JPA, KSM).

## Acknowledgements

We thank Dr. Sung Hyun Cho at the Penn State Huck Cryo-Electron Microscopy Facility for assistance with cryo-EM grid preparation and operating microscope for cryo-EM grid screening and cryo-EM data collection, and we thank Dr. Tatiana Laremore at Penn State Huck Proteomics and Mass Spectrometry Facility for technical assistance in protein identification. We thank Olivia Abboud for *D. melanogaster* maintenance. Research reported in this publication was supported by the Office of the Director, NIH, under award number S10OD026822-01 (S.L.H.). The results generated in this paper were computed on a hardware generously provided to us through the NVIDIA Academic Grant by NVIDIA Corporation. Figure 1A was created with BioRender.com.

## Data and materials availability

All data needed to evaluate the conclusions of this paper are present in the Main Text and/or the Supplementary Materials. The cryo-EM maps and the refined models were deposited in the Electron Microscopy DataBank (EMDB, www.ebi.ac.uk/emdb/) and Protein Data Bank (PDB, www.rcsb.org), respectively. Pol II EC 2.86 A overall: EMD-48624; Pol II-EC with stalk: EMD-48622 and 9MU7; Pol II EC without stalk: EMD-48623 and 9MU8; nucleosome: EMD-48619 and 9MU4; hexasome: EMD-48620 and 9MU5; nucleosome EC: EMD-48625 (overall), EMD-48626 and 9MU9 (composite); DLST: EMD-48621 and 9MU6.

## Supporting information

This article contains supporting information.

## Supplementary Figure Legends

**SI Appendix Figure 1.**
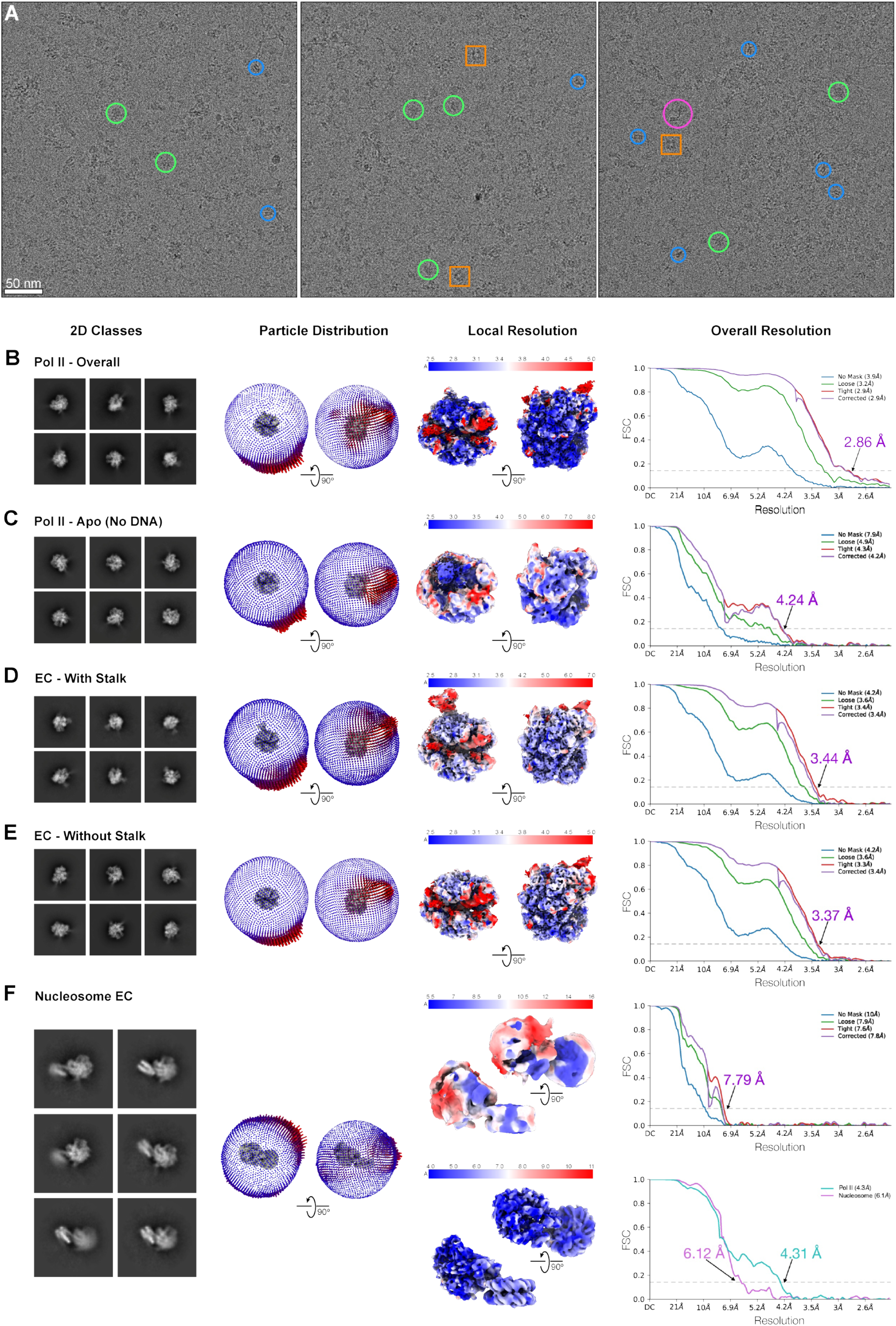
Cryo-EM micrograph and analysis. **(A)** Representative cryo-EM micrographs showing heterogeneity of the affinity-purified Pol II complex. Examples of different types of particles are marked: polymerases (green circle), nucleosomes (blue circles), nucleosome EC (pink circle), DLST (orange square) **(B-F)** Statistics on final reconstructions of apo-form Pol II (**B**) and ECs (**C-F**) including selected 2D class averages (left), Euler angle distribution of particles used in final reconstructions (second to left), local resolution maps with color ranging from blue to red representing high to low resolutions (second from right), and FSC curves calculated in CryoSPARC reported at the 0.143 FSC cutoff (right). In the Euler angle plot, the length of the cylinders is correlated to the distribution of particles in that direction. In the local resolution visualization, the maps are colored locally according to the scale above.

**SI Appendix Figure 2.**
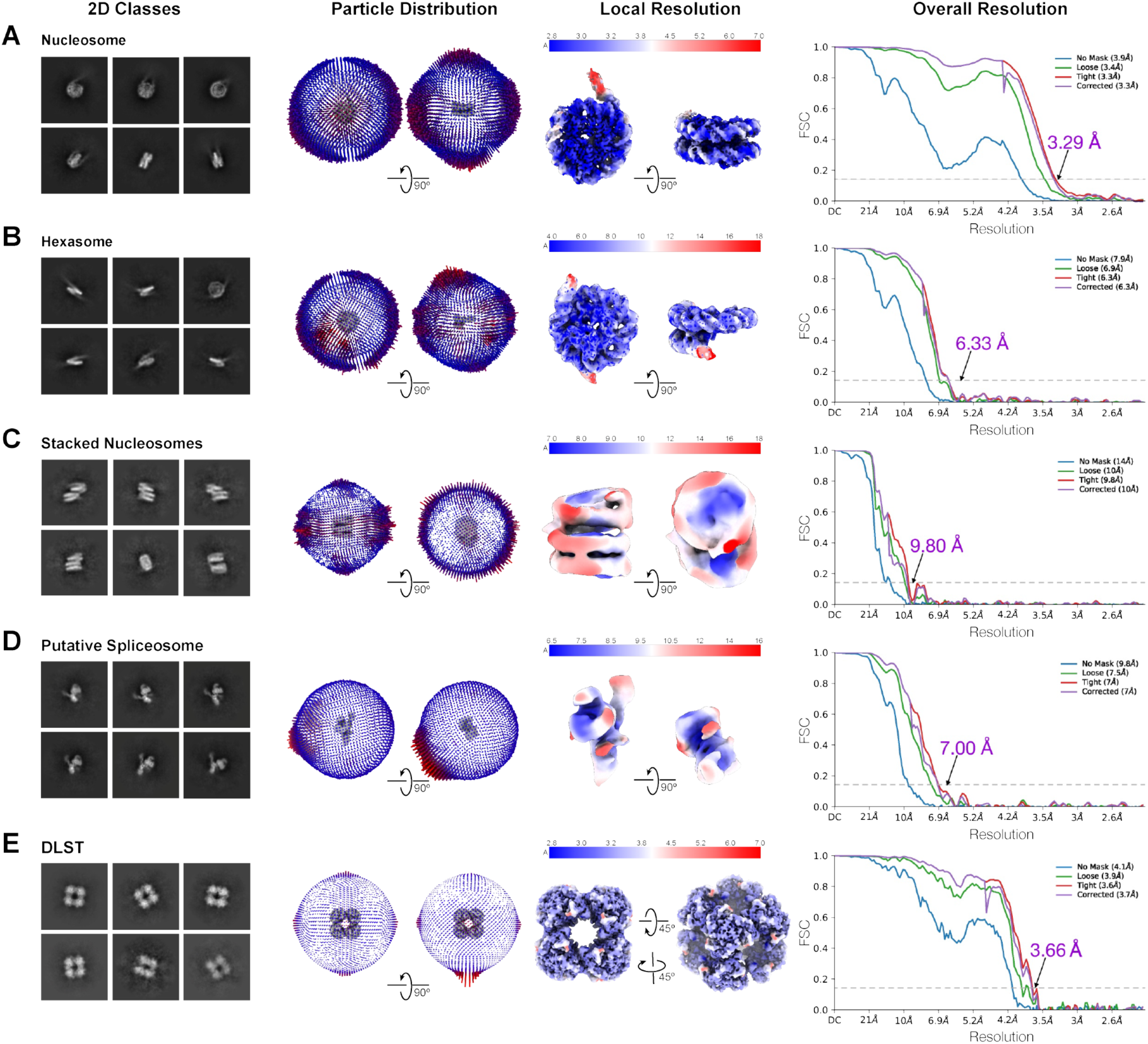
Cryo-EM micrograph and analysis, continued. **(A-E)** Statistics on final reconstructions of nucleosome (**A**), hexasome (**B**), di-nucleosomes (**C**), putative spliceosome (**D**) and DLST (**E**) including selected 2D class averages (left), Euler angle distribution of particles used in final reconstructions (second to left), local resolution maps with color ranging from blue to red representing high to low resolutions (second from right), and FSC curves calculated in CryoSPARC reported at the 0.143 FSC cutoff (right). In the Euler angle plot, the length of the cylinders is correlated to the distribution of particles in that direction. In the local resolution visualization, the maps are colored locally according to the scale above.

**SI Appendix Figure 3.**
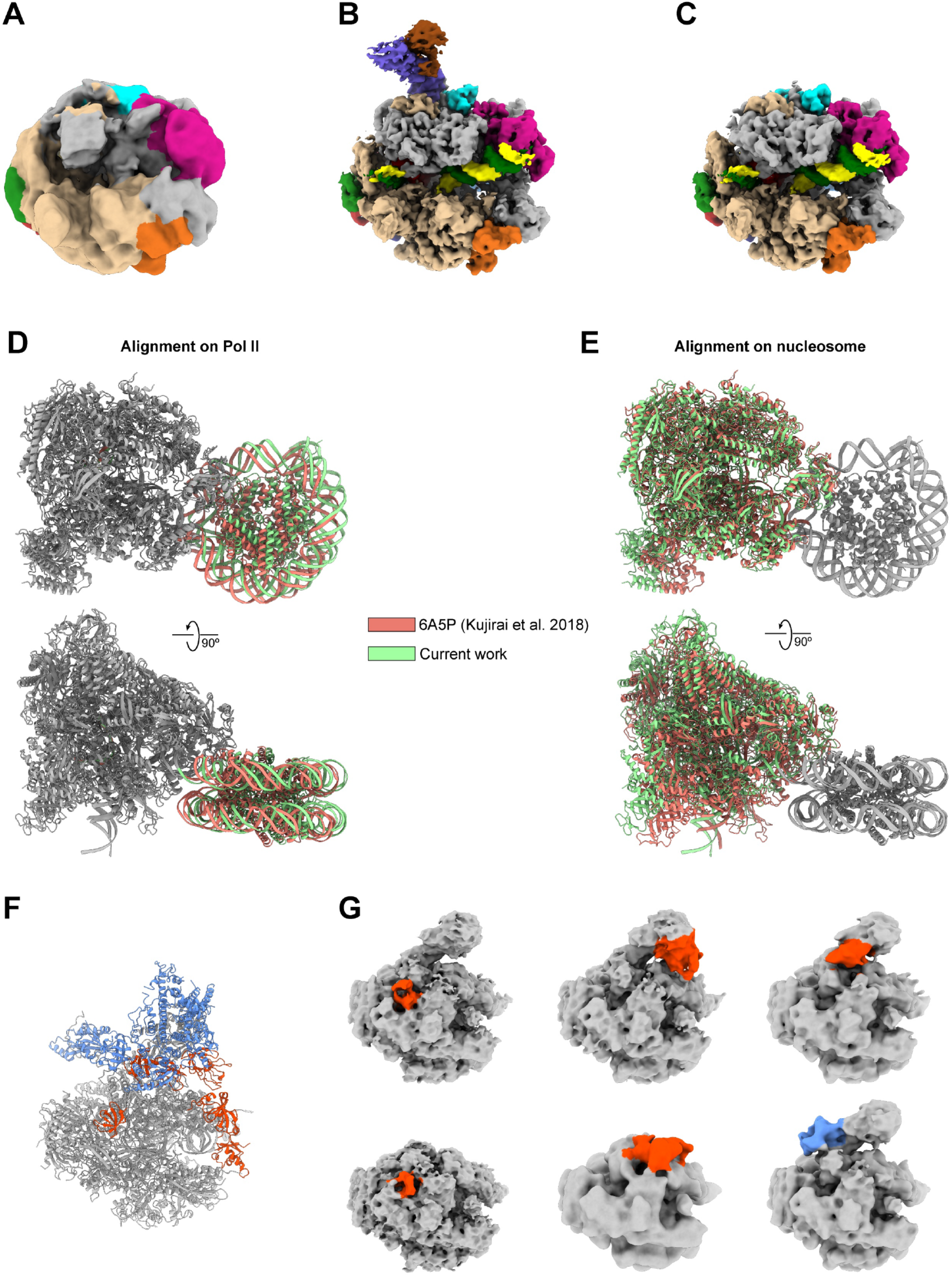
Variability of cryo-EM densities of native EC. **(A-C)** Cryo-EM densities of the apo-form Pol II (**A**), EC with the stalk **(B)** and EC without the stalk **(C)** with each subunit colored as described in **Figure 3. (D-E)** Comparison of nucleosome ECs from *Drosophila* (green, this study) and yeast paused at SHL(−5) (red, PDB: 6A5P) (8) with structures aligned on Pol II **(D)** and the nucleosome **(E)**. **(F)** Ribbon model of mammalian Pol II with elongation factor DSIF in red and histone chaperone SPT6 in blue (PBD: 6GMH (16)). **(G)** 3D classification of the Drosophila Pol II EC data focused on the DSIF or Spt6 binding sites reveal low resolution densities showing that some binding sites are occupied (red, DSIF; blue, Spt6), which may indicate co-purification of DSIF and Spt6 with endogenous Pol II. Mass spectrometry detected DSIF and Spt6 in high abundance.

**SI Appendix Figure 4.**
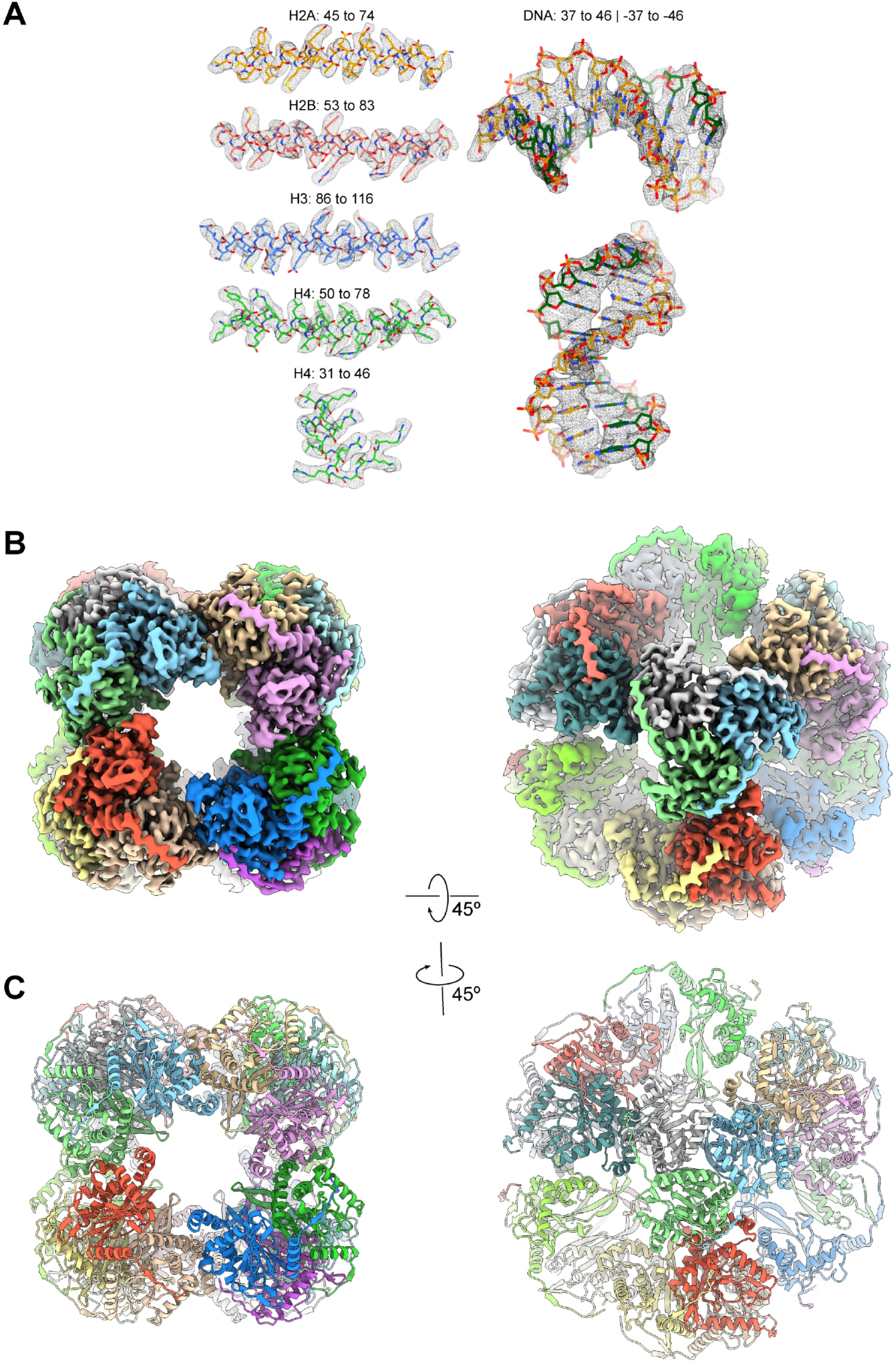
Quality of cryo-EM of nucleosome reconstruction; Cryo-EM map and model of DLST. **(A)** Selected representative 3.29 A resolution cryo-EM densities of core histones and DNA in nucleosome superimposed on the refined models. **(B)** Two views of the cryo-EM density of *Drosophila* dihydrolipoyllysine-residue succinyltransferase component of mitochondrial 2-oxoglutarate dehydrogenase complex at 3.66 A resolution, with individual subunits colored separately. **(C)** Two views of the refined model of DLST with individual subunits colored separately.

**SI Appendix Dataset 1. Protein identification by mass spectrometry.**

**SI Appendix Movie 1. Cryo-EM density map of EC with and without Rpb4/7 stalk.** Overview of the cryo-EM maps of the EC with and without the stalk. Related to Fig. 3A

**SI Appendix Movie 2. Cryo-EM density map of nucleosome EC.** Overview of the cryo-EM composite map of the nucleosome EC. Related to Fig. 5.

## Supplementary Info Appendix

**SI Appendix Table 1.**
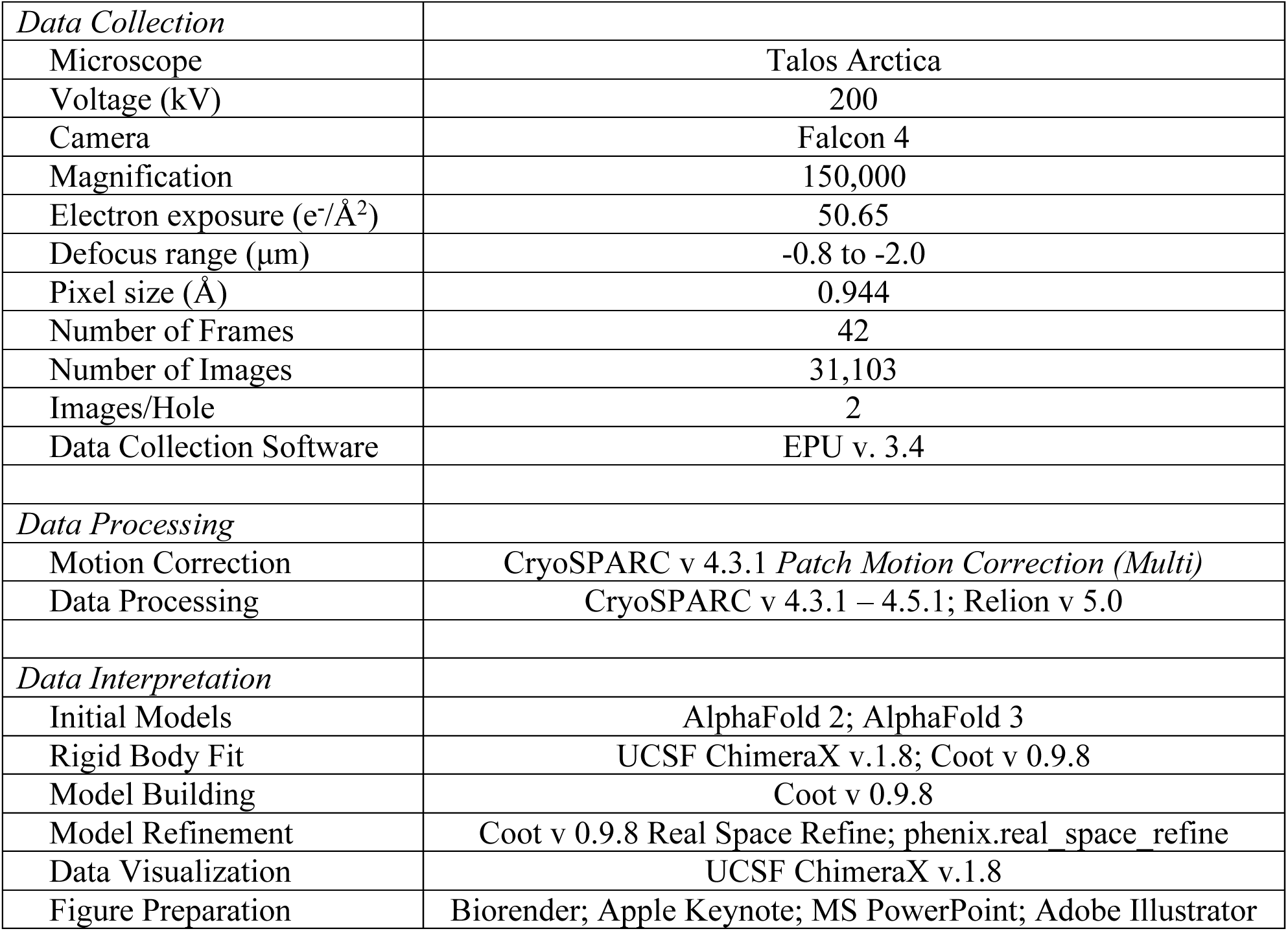
Cryo-EM data collection, interpretation and visualization details.

|  |  |
| --- | --- |
| *Data Collection* |  |
| Microscope | Talos Arctica |
| Voltage (kV) | 200 |
| Camera | Falcon 4 |
| Magnification | 150,000 |
| Electron exposure (e<sup>-</sup>/Å<sup>2</sup>) | 50.65 |
| Defocus range (μm) | -0.8 to -2.0 |
| Pixel size (Å) | 0.944 |
| Number of Frames | 42 |
| Number of Images | 31,103 |
| Images/Hole | 2 |
| Data Collection Software | EPU v. 3.4 |
| *Data Processing* |  |
| Motion Correction | CryoSPARC v 4.3.1 *Patch Motion Correction (Multi)* |
| Data Processing | CryoSPARC v 4.3.1 – 4.5.1; Relion v 5.0 |
| *Data Interpretation* |  |
| Initial Models | AlphaFold 2; AlphaFold 3 |
| Rigid Body Fit | UCSF ChimeraX v.1.8; Coot v 0.9.8 |
| Model Building | Coot v 0.9.8 |
| Model Refinement | Coot v 0.9.8 Real Space Refine; phenix.real_space_refine |
| Data Visualization | UCSF ChimeraX v.1.8 |
| Figure Preparation | Biorender; Apple Keynote; MS PowerPoint; Adobe Illustrator |

**SI Appendix Table 2.** Data processing and model refinement statistics for EC and Nucleosome EC.

| **Refinements** | **Pol II overall**<br>(EMD-48624) | **EC with stalk**<br>(EMD-48622)<br>(PDB: 9MU7) | **EC without stalk**<br>(EMD-48623)<br>(PDB: 9MU8) | **Nucleosome EC\*\***<br>(EMD-48625)<br>(EMD-48626)<br>(PDB: 9MU9) |
| --- | --- | --- | --- | --- |
| Initial # of particles | 876,764 | 876,764 | 876,764 | 36,552 |
| Final # of particles | 68,716 | 70,116 | 96,409 | 36,552 |
| Map resolution (Å) at 0.143 FSC threshold | 2.86 | 3.44 | 3.37 | 7.79/4.31/6.12* |
| Reference models (PDB code) | - | 6GML | 6GML | 6GML/3LZ0 |
| *Model composition* |  |  |  |  |
| Non-hydrogen atoms | - | 33,887 | 31,485 | 42,423 |
| Protein residues | - | 3,992 | 3,689 | 4,446 |
| Nucleic acid residues | - | 99 | 99 | 347 |
| Ligands | - | Zn:8 | Zn:8 | Zn:8 |
| *B factors (Å<sup>2</sup>)* |  |  |  |  |
| Protein | - | 13.6/491.5/117.2 | 57.2/403.6/112.2 | 30.0/1229.4/457.3 |
| DNA/RNA | - | 39.0/536.2/171.1 | 39.0/536.2/171.1 | 200.1/1104.7/352.9 |
| Ligand | - | 127.7/181.2/156.5 | 127.7/181.2/156.5 | 286.1/944.0/570.4 |
| *r.m.s. deviations* |  |  |  |  |
| Bond lengths (Å) | - | 0.006 | 0.006 | 0.003 |
| Bond angles (°) | - | 0.754 | 0.746 | 0.728 |
| *Validation* |  |  |  |  |
| MolProbity score | - | 1.48 | 1.48 | 1.48 |
| Clash score | - | 4.54 | 4.57 | 5.07 |
| Q-score | - | 0.393 | 0.403 | 0.202 |
| Rotamer outliers (%) | - | 0.00 | 0.00 | 0.00 |
| *Ramachandran plot* |  |  |  |  |
| Favored (%) | - | 96.29 | 96.26 | 96.61 |
| Allowed (%) | - | 3.71 | 3.74 | 3.39 |
| Disallowed (%) | - | 0.00 | 0.00 | 0.02 |

\* left, overall resolution; middle, focused on Pol II; right, focused on nucleosome
\*\* validation performed against the nucleosome EC composite map

**SI Appendix Table 3.** Data processing and model refinement statistics for nucleosome, hexasome and DLST.

| **Refinements** | **Nucleosome**<br>(EMD-48619)<br>(PDB: 9MU4) | **Hexasome**<br>(EMD-48620)<br>(PDB: 9MU5) | **DLST**<br>(EMD-48621)<br>(PDB: 9MU6) |
| --- | --- | --- | --- |
| Initial # of particles | 1,848,096 | 231,425 | 70,106 |
| Final # of particles | 66,988 | 38,738 | 23,543 |
| Map resolution (Å) at 0.143 FSC threshold | 3.29 | 6.33 | 3.66 |
| Reference models (PDB code) | 3LZ0 | Octameric nucleosome | AlphaFold |
| *Model composition* |  |  |  |
| Non-hydrogen atoms | 12,794 | 9,666 | 43,134 |
| Protein residues | 770 | 532 | 5,616 |
| Nucleic acid residues | 328 | 266 | - |
| Ligands | - | - | - |
| *B factors (Å<sup>2</sup>) (mean)* |  |  |  |
| Protein | 23.6/168.3/67.5 | 124.0/1398.6/616.9 | 40.6/144.90/78.1 |
| DNA | 15.7/472.4/123.3 | 188.7/1043.1/444.0 | - |
| Ligand | - | - | - |
| *r.m.s. deviations* |  |  |  |
| Bond lengths (Å) | 0.006 | 0.004 | 0.005 |
| Bond angles (°) | 0.837 | 0.722 | 0.983 |
| *Validation* |  |  |  |
| MolProbity score | 1.28 | 1.45 | 1.86 |
| Clash score | 2.50 | 5.89 | 7.51 |
| Q-score | 0.442 | 0.206 | 0.39 |
| Rotamer outliers (%) | 0.00 | 0.00 | 0.00 |
| *Ramachandran plot* |  |  |  |
| Favored (%) | 96.29 | 97.31 | 93.09 |
| Allowed (%) | 3.71 | 2.69 | 6.91 |
| Disallowed (%) | 0.00 | 0.00 | 0.00 |

**SI Appendix Table 4.** Protein sequences used for model building.

| **Protein Name** | **Gene Name** | **Uniprot ID** |
| --- | --- | --- |
| DNA-directed RNA polymerase II subunit RPB1 | RPB1 | P04052 |
| DNA-directed RNA polymerase II subunit RPB2 | RPB2 | P08266 |
| DNA-directed RNA polymerase II subunit RPB3 | RPB3 | O97183 |
| DNA-directed RNA polymerase II subunit RPB4 | RPB4 | Q9VEA5 |
| DNA-directed RNA polymerase II subunit RPB5 | RPB5 | Q7JZF5 |
| DNA-directed RNA polymerase II subunit RPB6 | RPB6 | Q24320 |
| DNA-directed RNA polymerase II subunit RPB7 | RPB7 | Q9VFB5 |
| DNA-directed RNA polymerase II subunit RPB8 | RPB8 | Q9VNZ3 |
| DNA-directed RNA polymerase II subunit RPB9 | RPB9 | P36958 |
| DNA-directed RNA polymerase II subunit RPB10 | RPB10 | Q9VC49 |
| DNA-directed RNA polymerase II subunit RPB11 | RPB11 | Q9VJE4 |
| DNA-directed RNA polymerase II subunit RPB12 | RPB12 | Q6IGE3 |
| Histone H2A | H2A | P84051 |
| Histone H2B | H2B | P02283 |
| Histone H3 | H3 | P02299 |
| Histone H4 | H4 | P84040 |
| Dihydrolipoyllysine-residue succinyltransferase<br>component of 2-oxoglutarate dehydrogenase<br>complex, mitochondrial | DLST | Q9VGQ1 |

